# Anti-inflammatory effects of WFS1 in pancreatic β-cells

**DOI:** 10.1101/2022.02.09.479773

**Authors:** Shuntaro Morikawa, Lindsey Blacher, Chinyere Onwumere, Fumihiko Urano

## Abstract

Wolfram syndrome is a rare genetic disorder characterized by juvenile-onset diabetes mellitus, optic nerve atrophy, hearing loss, diabetes insipidus, and progressive neurodegeneration. Pathogenic variants in the *WFS1* gene are the main causes of Wolfram syndrome. *WFS1* encodes a transmembrane protein localized to the endoplasmic reticulum (ER) and regulates the unfolded protein response (UPR). Loss of function of *WFS1* leads to dysregulation of insulin production and secretion, ER calcium depletion, and cytosolic calpains activation, resulting in activation of apoptotic cascades. Although the terminal UPR has been shown to induce inflammation that accelerates β-cell dysfunction and death in diabetes, the contribution of β-cell inflammation to the development of diabetes in Wolfram syndrome has not been fully understood. Here we show that *WFS1*-deficiency enhances the gene expression of inflammatory cytokines and chemokines, leading to cytokine-induced ER-stress and cell death in pancreatic β-cells. PERK and IRE1α pathways mediate high glucose-induced inflammation in a β-cell model of Wolfram syndrome. M1-macrophage infiltration and hypervascularization are seen in the pancreatic islets of *Wfs1* whole-body knockout mice, demonstrating that WFS1 regulates anti-inflammatory responses in β-cells. Our results demonstrate that inflammation plays an essential role in the progression of β-cell death and diabetes in Wolfram syndrome and reveal that the pathways involved in ER stress-mediated inflammation provide potential therapeutic targets for the treatment of Wolfram syndrome.

## Introduction

Wolfram syndrome is a rare genetic disorder characterized by juvenile-onset diabetes mellitus, optic nerve atrophy, hearing loss, and progressive neurodegeneration (1-3). Most cases of Wolfram syndrome are caused by pathogenic variants in the *WFS1* gene, which encodes a transmembrane protein localized to the endoplasmic reticulum (ER) (4). WFS1 regulates ER calcium homeostasis, and its dysfunction causes the accumulation of unfolded/misfolded proteins in the ER (referred to as ER stress). Pathological ER stress mediates cell death in pancreatic β-cells and neuronal cells, which is thought to be the mechanism of Wolfram syndrome development (5-7).

ER stress occurs even under physiological conditions, as protein folding in the ER is an error-prone process. To maintain cellular and organ homeostasis, the unfolded protein response (UPR) determines the cell’s fate by sensing physiological or pathological ER stress (8). The induction of adaptive UPR that leads to cell survival, or the initiation of pathological UPR that leads to cell apoptosis, is regulated by the molecular components of the UPR: inositol-requiring transmembrane kinase/endoribonuclease 1α (IRE1α), protein kinase RNA-like endoplasmic reticulum kinase (PERK), and activating transcription factor 6 (ATF6).

Increasing evidence indicates that pathological ER stress causes sterile inflammation, an inflammatory response regulated by the pathological UPR (9). Among the UPR signaling pathways, the IRE1α pathway is known to degrade IκB, a specific inhibitor of nuclear factor-κB (NF-κB), which is a family of transcription factors that regulates the gene expression of pro-inflammatory cytokines (10). In addition, the PERK pathway is known to suppress IκB translation (11). These UPR pathways regulate the nuclear translocation of NF-κB and are the molecular mechanisms linking pathogenic ER stress and sterile inflammation (11). Since β-cells are heavily loaded with protein synthesis and proinsulin is prone to misfold, ER stress is closely associated with the pathogenesis of diabetes (12). Moreover, increasing evidence suggests that ER stress-induced sterile inflammation is involved in the progression of type 2 diabetes (12-14).

We previously showed that WFS1 dysfunction causes pathological ER stress-mediated β-cell death (15). However, the role of inflammation in Wolfram syndrome and its relationship with WFS1 has not been fully elucidated. In this study, we hypothesized that pathogenic ER stress and inflammation in β-cells might accelerate the progression of diabetes in Wolfram syndrome. Our findings reveal islet-localized inflammation in Wolfram syndrome models and suggest that WFS1 functions to regulate sterile inflammation in β-cells. Supplementing our previous model that shows the relationship between WFS1 and ER stress-induced β-cell death, here we provide new evidence showing that islet-localized inflammation accelerates the progression of diabetes in Wolfram syndrome.

## Results

### Loss of function of WFS1 induces sterile inflammation in pancreatic β-cells

To investigate whether WFS1 dysfunction leads to sterile inflammation via pathological ER stress in β-cells, we first examined apoptosis and pro-inflammatory cytokine gene expression levels in *Wfs1* knockdown INS-1E cells. In this *in vitro* Wolfram syndrome model, cleaved caspase-3 levels were increased, suggesting activation of apoptosis (Figure 1A, 1B). Furthermore, the gene expression levels of pro-inflammatory cytokines (*Il-1β, Il-6*, and *Cxcl1*(*Il-8)*) and *Chop* were significantly upregulated (Figure 1C). These results indicate that WFS1 plays a role in the regulation of ER stress-mediated cell death and pro-inflammatory cytokine gene expression.

**Figure 1.**
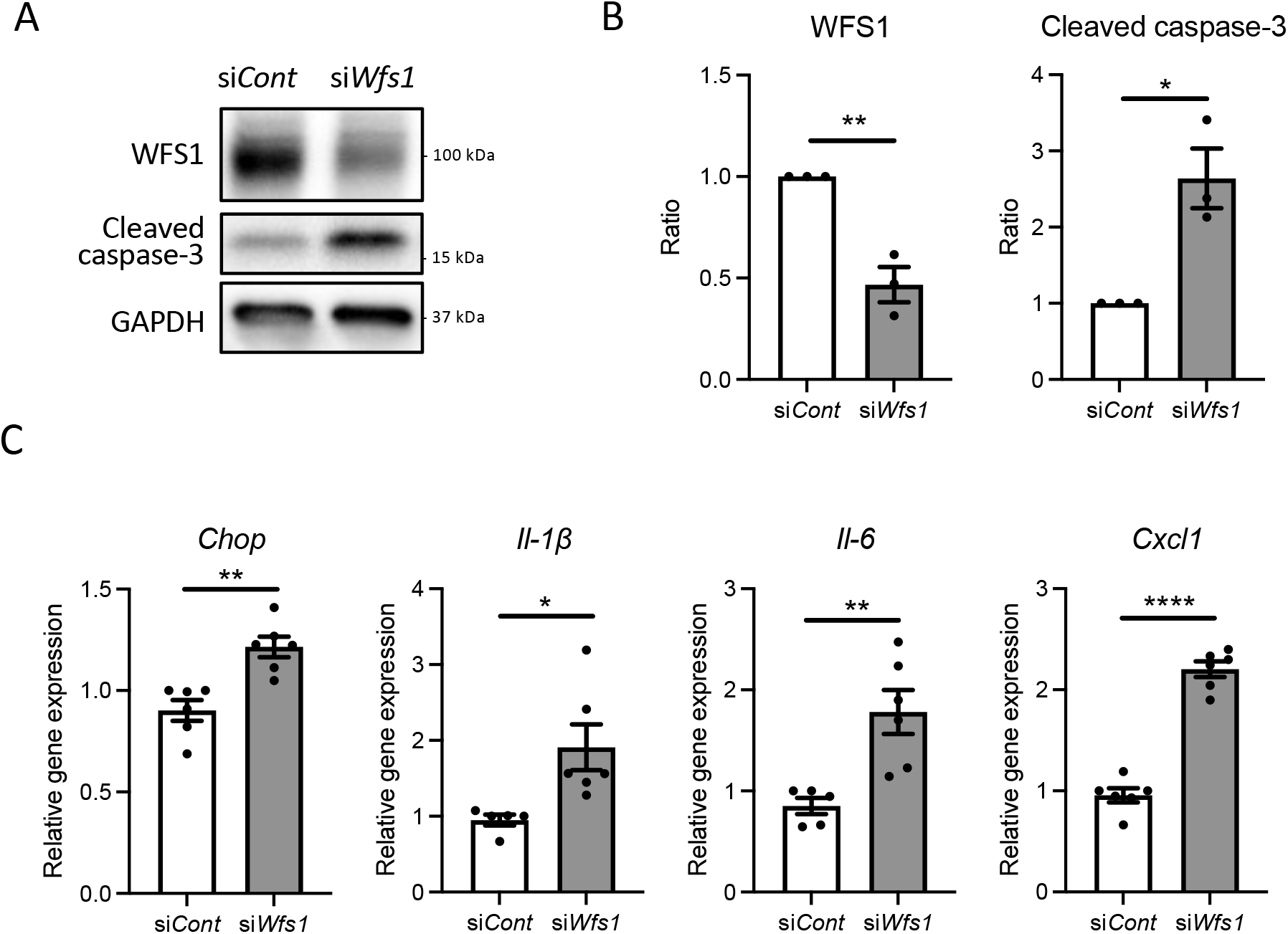
Cell apoptosis and expression of pro-inflammatory cytokine genes are upregulated in *Wfs1*-deficient pancreatic β-cells. (A, B) INS-1E cells were transfected with scrambled siRNA (si*Cont*) or siRNA targeting rat *Wfs1* (si*Wfs1*). WFS1 and cleaved caspase-3 protein expression levels were analyzed by immunoblot. Representative images are shown in (A), and quantitative analysis results are shown in (B) (n=3, normalized to GAPDH). (C) mRNA expression levels of *Chop, Il-1β, Il-6*, and *Cxcl1* (*Il-8*) in INS-1E cells treated with si*Cont* or si*Wfs1* normalized to *18srRNA* (n=5-6). Data are shown as mean ± SEM, * P<0.05, ** P<0.005, **** P<0.0001.

### Pro-inflammatory cytokine gene expression is enhanced by high-glucose stimulation in β-cell models of Wolfram syndrome

Diabetes, caused by ER stress-mediated pancreatic β-cell apoptosis, is one of the major and early manifestations of Wolfram syndrome (1, 16), suggesting that pancreatic β-cells of Wolfram syndrome patients are exposed to hyperglycemia for extended time periods. Therefore, we investigated how WFS1-deficient pancreatic β-cells produce pro-inflammatory cytokines under a chronic high-glucose environment. First, we examined the influence of chronic high-glucose conditions on cell death and ER stress using *Wfs1*-knockout INS-1 832/13 cells (*Wfs1*-KO INS-1 cells) (Figure 2A). These *Wfs1*-KO INS-1 cells possess the characteristics of pancreatic β-cells in Wolfram syndrome: thapsigargin intolerance and mitochondrial dysfunction (17) (Figure S1). *Wfs1*-KO INS-1 cells treated with high glucose showed elevated proinsulin expression, caspase-3/7 activity, and gene expression levels of ER stress markers such as *Chop, sXbp1, Bip*, and *Txnip* compared to wild type INS-1 832/13 cells (*Wfs1*-WT INS-1 cells), (Figure 2B, 2C, 2D). In this condition, the gene expression levels of pro-inflammatory cytokines (*Il-1β, Il-6*), chemokine (*Ccl2*), and vascular endothelial growth factor A (*VegfA*) were increased in *Wfs1*-KO INS-1 cells (Figure 2E). The chronic high-glucose condition enhanced not only the cell apoptosis but also the expression of pro-inflammatory cytokine genes in *Wfs1*-KO INS-1 cells.

**Figure 2.**
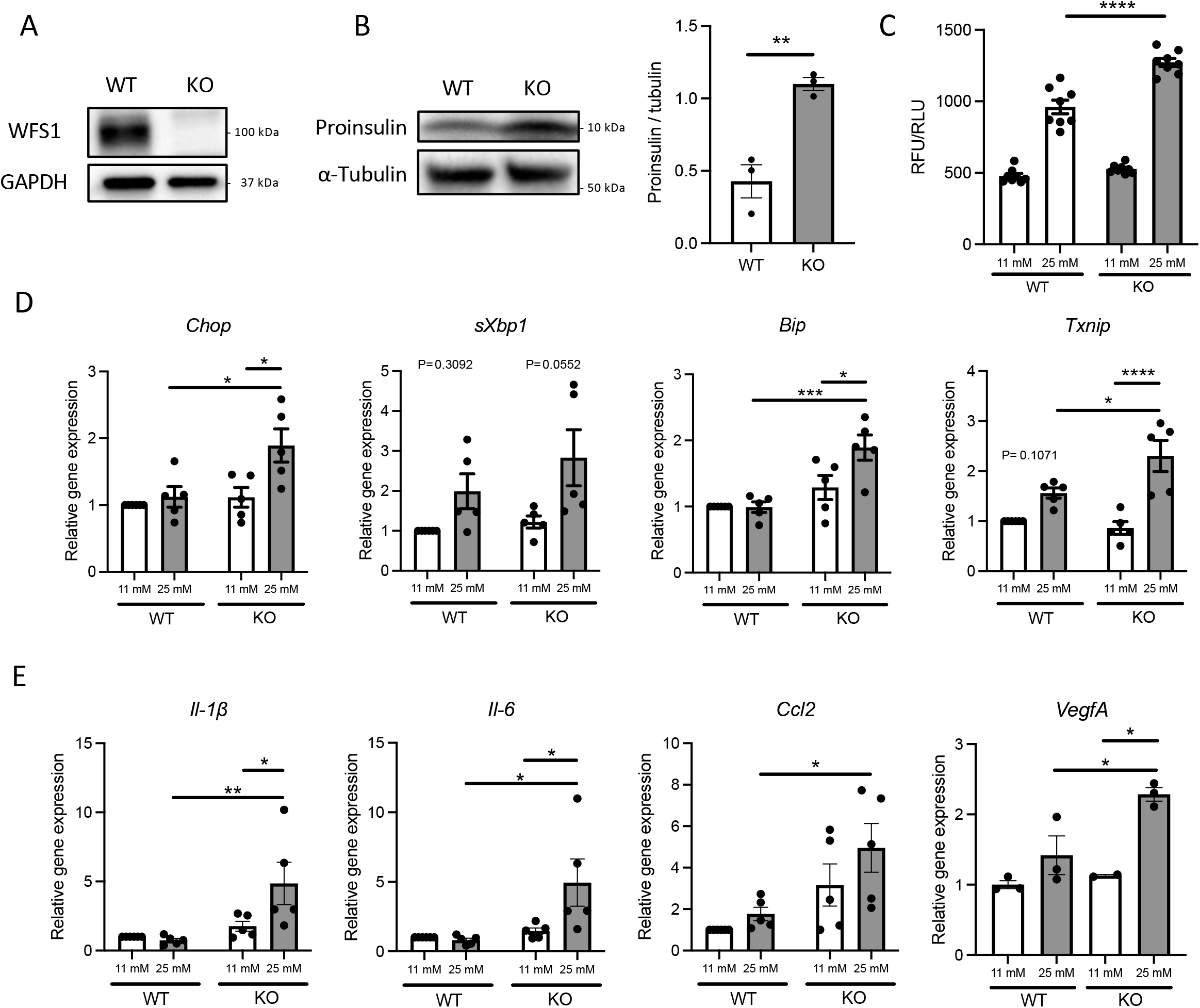
Hyperglycemia upregulates pro-inflammatory cytokine genes expression in *Wfs1*-deficient pancreatic β-cells. (A) Immunoblot images of WFS1 in *Wfs1* wild type (WT) and knock-out (KO) INS-1 832/13 cells. (B) Left panel: Immunoblot images of proinsulin in WT and KO INS-1 832/13 cells treated with 25 mM glucose for 1 h. Right panel: Proinsulin band intensity was quantified and normalized to tubulin (n=3). (C) Caspase-3/7 activity normalized to cell viability in WT and KO INS-1 832/13 cells treated with 11 mM or 25 mM glucose for 48 h (n=8). (D) mRNA expression levels of *Chop, sXbp1, Bip*, and *Txnip* in WT or KO INS-1 832/13 cells treated with 11 mM or 25 mM glucose for 48 h (n=3-5), normalized to *18srRNA*. (E) The mRNA expression level of *Il-1β, Il-6, Ccl2*, and *VegfA* in WT and KO INS-1 832/13 cells treated with 11 mM or 25 mM glucose for 48 h (n=3-5), normalized to *18srRNA*. Data are shown as mean ± SEM, * P<0.05, ** P<0.005, *** P<0.001, **** P<0.0001.

### Cytokine treatment enhances ER stress-mediated cell apoptosis and pro-inflammatory cytokine gene expression in β-cell models of Wolfram syndrome

Pro-inflammatory cytokines secreted from pancreatic β-cells are known to act in a paracrine or autocrine manner in type 2 diabetes models (18). Therefore, we hypothesized that the locally secreted cytokines might influence the characteristics of pancreatic β-cells in Wolfram syndrome. To test this idea, we evaluated the cell death and ER stress markers in *Wfs1*-KO INS-1 cells treated with cytokines. *Wfs1*-KO INS-1 treated with IFN-γ and IL-1β showed the enhanced ER stress-mediated cell death (Figure 3A). Furthermore, the gene expression levels of *Il-1β, Il-6*, and *Ccl2* were higher in *Wfs1*-KO INS-1 cells compared to *Wfs1*-WT INS-1 cells (Figure 3B). These results indicate that cytokine treatment enhances ER stress-mediated cell death and upregulates the pro-inflammatory cytokine gene expression in *Wfs1*-deficient β-cells.

**Figure 3.**
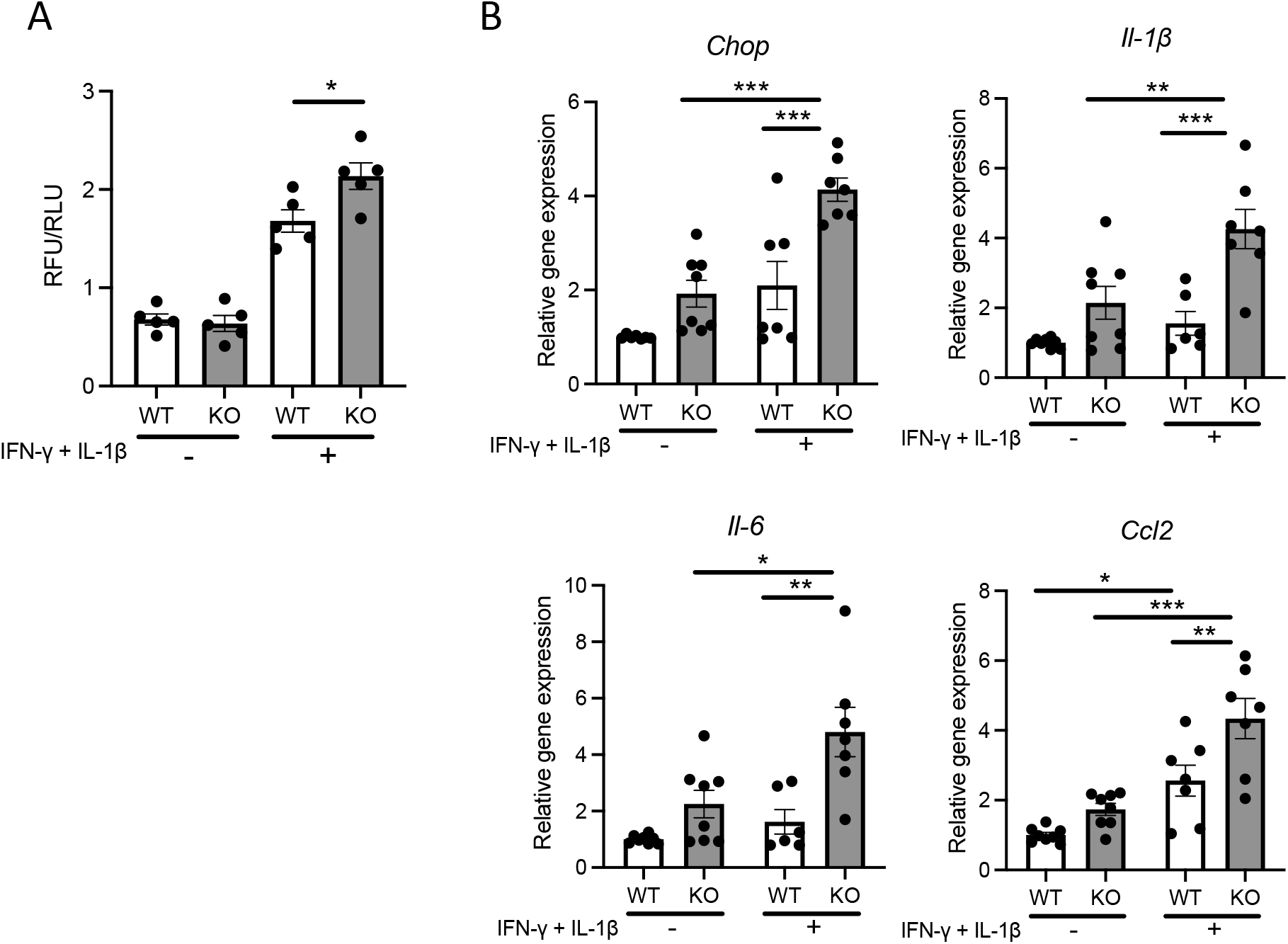
Cytokine treatment causes ER-stress induced cell death and pro-inflammatory cytokines gene upregulation in *Wfs1*-deficient pancreatic β-cells. (A) Caspase-3/7 activity normalized to cell viability in *Wfs1* wild type (WT) and knock-out (KO) INS-1 832/13 cells treated with IFN-γ (50 ng/mL) and IL-1β (50 ng/mL) for 24 h (n=5). (B) mRNA expression level of *Chop, Il-1β, Il-6*, and *Ccl2* in WT and KO INS-1 832/13 cells treated with IFN-γ (50 ng/mL) and IL-1β (50 ng/mL) for 24 h (n=6-8), normalized to *18srRNA*. Data are shown as mean ± SEM, * P<0.05, ** P<0.005, *** P<0.001.

### High-glucose induced pro-inflammatory cytokine gene expression is mediated by PERK pathway in Wolfram syndrome pancreatic β-cells

Next, we investigated the mechanisms underlying the upregulation of pro-inflammatory cytokine gene expressions in *Wfs1*-KO INS-1 cells. We first examined the protein expression level of nuclear-localized NF-κB, which is known to regulate the expression levels of pro-inflammatory cytokines genes (10). We observed enhanced NF-κB nuclear translocation in *Wfs1*-KO INS-1 cells following high-glucose treatment (Figure 4A).

**Figure 4.**
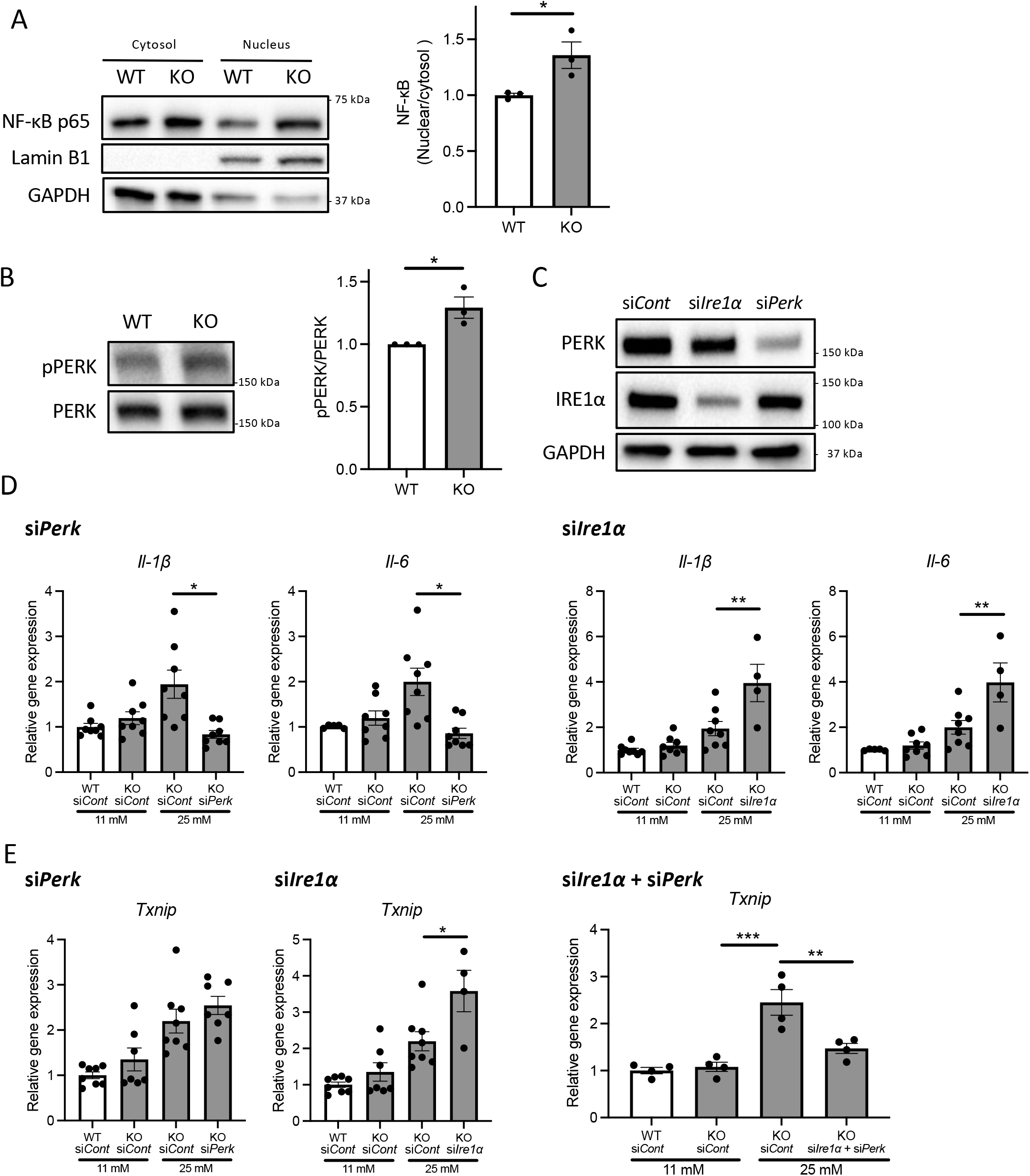
PERK pathway regulates the high-glucose induced pro-inflammatory gene expression in *Wfs1*-deficient pancreatic β-cells. (A) Left panel: Immunoblot images of NF-κB p-65 in the cytosolic or nuclear fraction of *Wfs1* wild type (WT) and knock-out (KO) INS-1 832/13 cells. WT and KO INS-1 832/13 cells were treated with 5 mM glucose for 18 h and then with 25 mM glucose for 30 min. Right panel: The ratio of nuclear and cytosolic NF-κB p-65 protein levels (n=3). (B) Left panel: Immunoblot images of pPERK and PERK. WT and KO INS-1 832/13 cells were treated with 5 mM for 16 h and then with 25 mM glucose for 24 h. Right panel: pPERK band intensity was quantified and normalized to total PERK (n=3). (C) Immunoblot images of PERK and IRE1α in WT and KO INS-1 832/13 cells treated with siRNA against *Perk* or *Ire1*α. (D) mRNA expression level of *Il-1β* and *Il-6* normalized to *18srRNA* in WT or KO INS-1 832/13 cells treated with 11 mM or 25 mM glucose for 30 h together with siRNA against *Perk* or *Ire1*α (n=4-8) (E) mRNA expression levels of *Txnip* normalized to *18srRNA* in WT and KO INS-1 832/13 cells treated with 11 mM or 25 mM glucose for 6 h together with siRNA against *Ire1*α and *Perk* (n=4). Data are shown in mean ± SEM, * P<0.05, ** P<0.005, *** P<0.001.

We next examined how high-glucose treatment activates the NF-κB pathway. As described above, the downstream pathways of pathological UPR were activated in high-glucose treated *Wfs1*-KO INS-1 cells, enhancing expression level of ER stress markers such as *Txnip*. TXNIP is known to activate the NLRP3 inflammasome, which leads to caspase-1 activation and IL-1β maturation, and its gene expression is regulated by PERK and IRE1 (19-21). Therefore, to determine whether the pathogenic UPR pathway enhances the NF-κB signaling pathways, we downregulated PERK and IRE1α in *Wfs1*-KO INS-1 cells. We found that the PERK was activated at baseline in high-glucose treated *Wfs1*-KO INS-1 cells (Figure 4B). Unlike knocking down IRE1α, knocking down PERK suppressed the enhanced expression of pro-inflammatory cytokine genes in high-glucose treated *Wfs1*-KO INS-1 cells (Figure 4C, 4D). These results suggest that the induction of pro-inflammatory cytokine gene expressions in high-glucose treated *Wfs1*-KO INS-1 cells was mainly mediated by the PERK pathway and subsequent NF-κB nuclear translocation. The increased *Txnip* expression in high-glucose treated *Wfs1*-KO INS-1 cells was suppressed by simultaneously knocking down PERK and IRE1α (Figure 4E). These results suggest that high-glucose treatment of *Wfs1*-KO-INS-1 cells enhances NF-κB-mediated pro-inflammatory cytokine genes expression via the PERK pathway and enhances TXNIP-mediated IL-1β processing through the PERK and IRE1α pathways.

### Islet-localized and systematic inflammation in the Wolfram syndrome mouse model

To confirm the findings above, we next investigated whether inflammation occurs in a mouse model of Wolfram syndrome. Although several WS mouse models have been developed, we used the *Wfs1* whole-body knockout 129S6 mice (*Wfs1*-KO mice) in this study (15, 22), which recapitulates human Wolfram syndrome phenotypes and progressively develops glucose tolerance impairment between 4.5 and 6.5 weeks old that persists through at least 10 months of age (15) (Figure S2A, S2B). Based on this knowledge, we collected the pancreatic tissues from mature-adult (16-week-old) and middle-aged (10-month-old) wild type (WT) and *Wfs1*-KO mice and investigated whether macrophages infiltrate the islets. Pancreatic tissue sections from each age were stained with Iba1, a marker for M1/M2 macrophage. Although there were no significant differences in 16-week-old mice, higher Iba1 density was observed in *Wfs1*-KO mice islets at 10 months age-old (Figure 5A, 5B). To further evaluate the character of these islet-resident macrophages, we performed imaging mass cytometry (IMC) on the *Wfs1*-KO mice pancreatic sections. The intra-islet macrophages in *Wfs1*-KO mice pancreatic islets expressed CD68 and IL-1β, indicating inflammatory M1-like characteristics (Figure 5C). There was also strong fibrosis in the islets of *Wfs1*-KO mice (Figure 5D, 5E), previously described as the hallmark of chronic inflammation in the diabetic islets (23). Together, these findings indicate the presence of inflammation in the *Wfs1*-KO mice islets.

**Figure 5.**
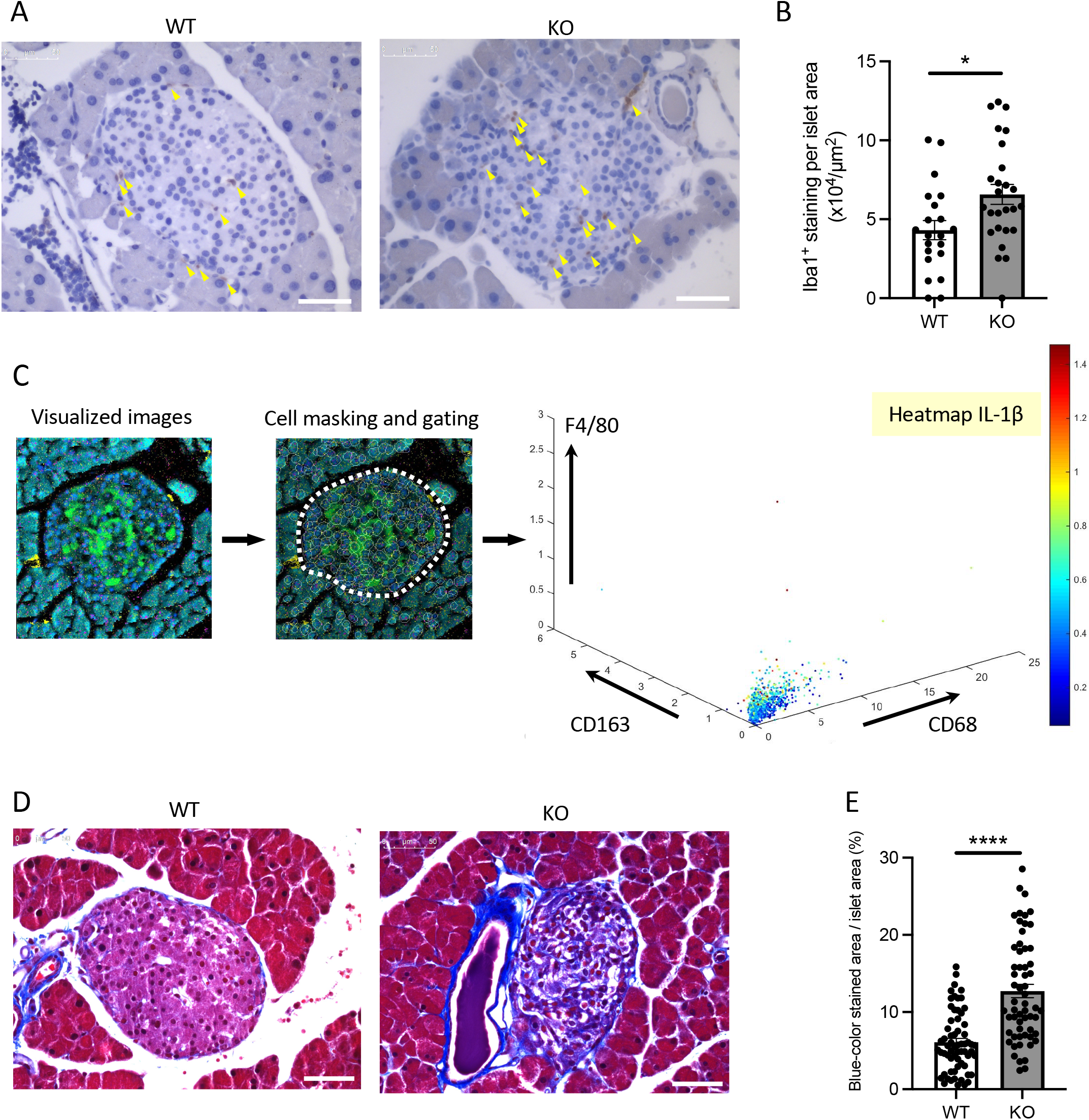
Macrophage infiltration and fibrosis in the Wolfram syndrome mouse model islets. (A) Iba1 DAB staining of the islets from *Wfs1* wild type (WT) and whole-body knockout 129S6 (KO) male mice at 10 months of age. Yellow arrowheads point to the stained macrophages. Scale bars 50 µm. (B) The number of Iba1 positive cell was normalized to each islet’s area size (WT, n=21; KO, n=28). (C) Left panel: Visualized imaging mass cytometry (IMC) image of KO male mouse stained with F4/80 (red), insulin (green), DNA (blue), CD68 (magenta), and CD163 (yellow). Contrast staining was performed with Ruthenium (cyan). Middle panel: cells were masked, and the islet area was gated. Right panel: Scatter plot of F4/80, CD163, and CD68. IL-1β was shown in the heatmap. (D) Trichrome-Masson staining of the islets from WT and KO male mice at 10 months of age. Collagen fibers are stained in blue. Scale bars 50 µm. (E) The blue-colored area was normalized to each islet’s area size (WT, n=33; KO, n=34). Data are shown in mean ± SEM, * P<0.05, **** P<0.001.

To examine whether the inflammation was also present in other tissues, we assessed cytokine levels in serum and bone marrow-derived macrophages of *Wfs1*-KO mice. Serum cytokine levels, notably interferon gamma-induced protein 10 (IP-10, CXCL10) was elevated in *Wfs1*-KO mice (Figure S3). These results suggest that the inflammation in these *Wfs1*-KO mice was not limited to the pancreatic islets. Although the high-glucose condition did not alter the gene expression levels of pro-inflammatory cytokines in bone marrow-derived macrophages (Figure S4A, S4B), advanced glycation end products (AGE) treatment enhanced *Ccl2* gene expression in bone marrow-derived macrophages of *Wfs1*-KO mice (Figure S4C, S4D).

### Hypervascularization in Wolfram syndrome mouse model islets

In *Wfs1*-KO INS-1 cells, we observed that high-glucose treatment enhanced *VegfA* gene expression (Figure 2E). VEGFA, an angiogenic factor, is expressed in the mouse islet and regulates vascular growth and permeability (24-26). Therefore, we tested if increased VEGFA expression may cause abnormal angiogenesis in the pancreatic islets of *Wfs1*-KO mice. We evaluated the endothelial cell marker (CD31) in the pancreas of WT and *Wfs1*-KO mice. A higher CD31 signal density was observed in *Wfs1*-KO mice islets compared to WT mice islets (Figure 6A, 6B). These results indicate the existence of hypervascularization in the islets of *Wfs1*-KO mice.

**Figure 6.**
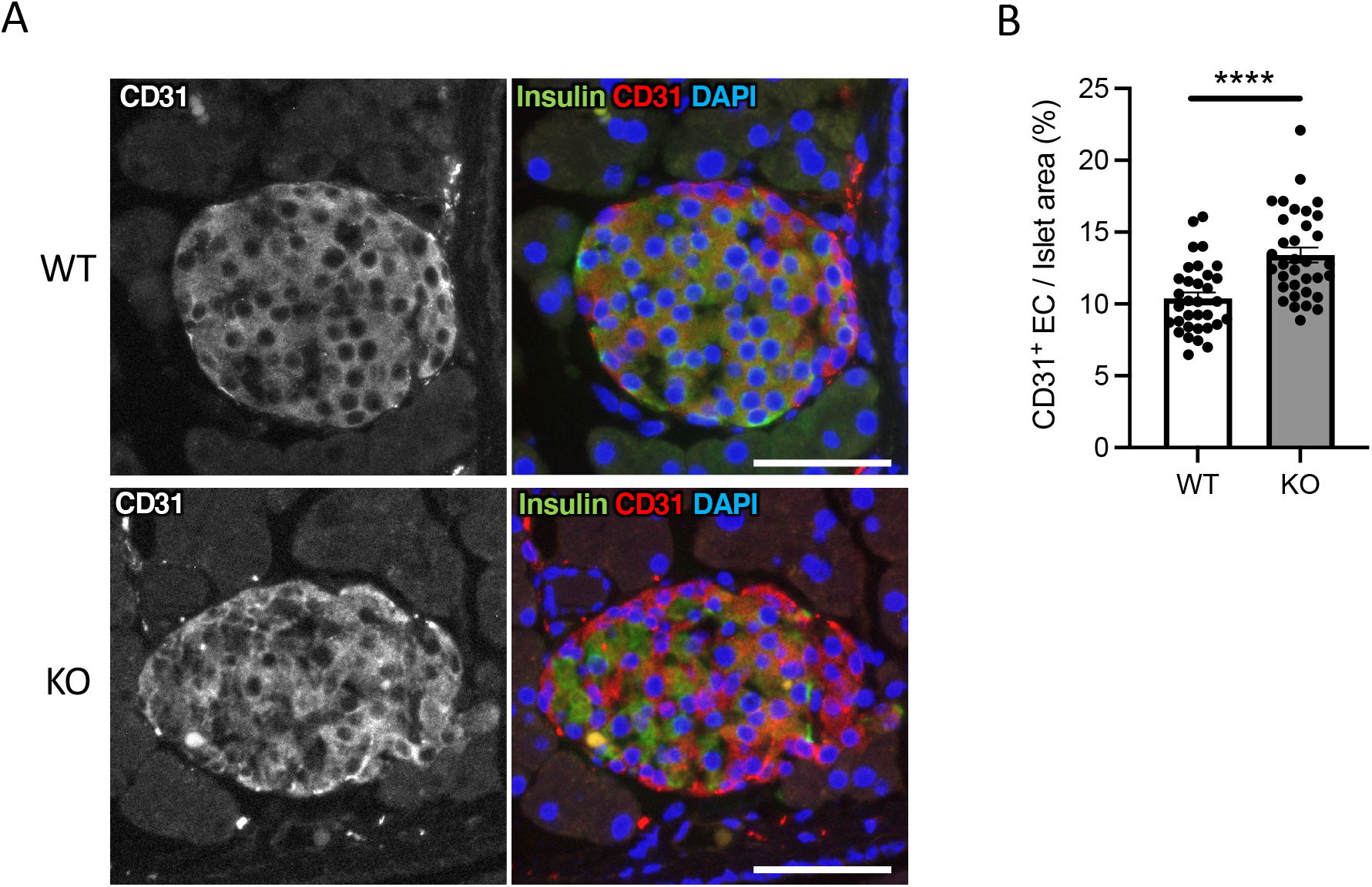
Hypervascularization in the islets Wolfram syndrome mouse model. (A) Representative immunofluorescent images of islets from *Wfs1* wild type (WT) and whole-body knockout 129S6 (KO) male mice at 10 months of age. Scale bars 50 µm. (B) Quantification of endothelial cell marker (CD31) positive area and islet area composition (WT, n=33; KO, n=34). Data are shown as mean ± SEM, **** P<0.001.

## Discussion

Although the relationship between the ER stress and pancreatic β-cell death in Wolfram syndrome has been well established, the role of inflammation in the development of Wolfram syndrome remains unknown. This study provides the first evidence indicating that high-glucose and cytokine treatment activate inflammation in β-cell and mouse models of Wolfram syndrome. We also confirmed the cell-nonautonomous pancreatic β-cell inflammation, including M1-macrophage infiltration in the islets of a mouse model of Wolfram syndrome. In pancreatic β-cells, ER stress and high-glucose conditions enhance the production and secretion of IL-1β, which is characteristic of sterile inflammation (19, 27). Our data demonstrate that the vulnerability to high-glucose and high-cytokine conditions potentiates ER stress and pro-inflammatory cytokine gene expression in WFS1-deficient pancreatic β-cells. This environment leads to pancreatic β-cell death, and the consequent chronic hyperglycemia induces sterile inflammation in WS, causing further pancreatic β-cell loss. This vicious cycle of inflammation and ER stress may accelerate the progression of diabetes in Wolfram syndrome (Figure 7).

**Figure 7.**
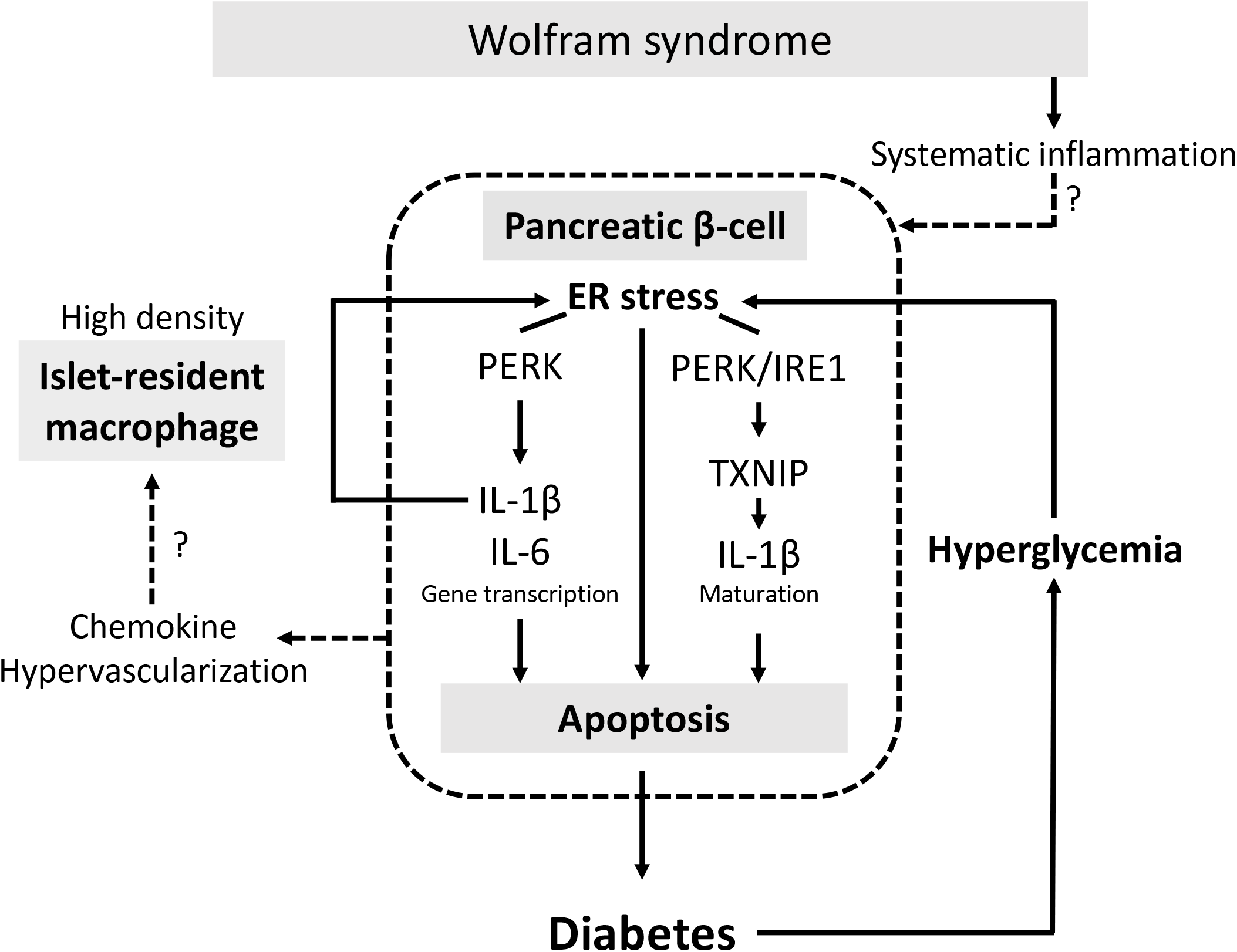
The vicious cycle of inflammation and ER stress accelerates the progression of diabetes in Wolfram syndrome. Schematic of the relationship between ER stress and sterile inflammation in Wolfram syndrome (WS). Diabetes in WS is caused by pathogenic ER stress-induced pancreatic β-cell apoptosis. Hyperglycemic conditions of *WFS1*-deficient pancreatic β-cells enhance pathogenic ER stress and induce sterile inflammation. Experimental high-glucose condition induces pro-inflammatory cytokine gene transcription via the PERK pathway and enhances TXNIP-mediated IL-1β processing through the PERK and IRE1α pathways. This high-cytokine condition accelerates ER stress-mediated cell death and upregulates the pro-inflammatory cytokine gene expression in *WFS1*-deficient pancreatic β-cells. Secreted chemokines or the hypervascularized environment in the WS islets may cause cell-nonautonomous pancreatic β-cell inflammation, including M1-macrophage infiltration.

The islet-resident macrophages originated from hematopoietic stem cells and are present at steady state conditions (28). However, the density of islet-resident macrophages is known to increase in patients with type 2 diabetes and rat and mouse models of type 2 diabetes (29, 30). These islet-resident macrophages express an M1-like transcript signature, and the macrophages infiltrated into the islets trigger subsequent autoimmune responses (30-32). In this study, we show that pancreatic islets in *Wfs1*-KO mice were highly infiltrated with inflammatory M1-macrophages. Therefore, islet-localized inflammation, which has been shown in the type 2 diabetes models, is also present in cell and mouse models of Wolfram syndrome.

Circulating macrophages are known to infiltrate tissues and differentiate into tissue-resident macrophages (33). There are two possible reasons for the increased density of islet-resident macrophages in the Wolfram syndrome mouse model. First, macrophages may respond to cytokines and chemokines that are thought to be secreted from islets, as shown by previous studies (30). Our results show that β-cell model of Wolfram syndrome under hyperglycemia expresses a higher level of *Ccl2* mRNA compared to the wild type cells. Furthermore, previous studies have reported that the recruitment of circulating macrophages to the islets is dependent on CCL2/CCR2 (29, 34). The second possible reason for the high macrophage infiltration is the increased angiogenesis observed in the Wolfram syndrome model mice islets. In this study, we show that *VegfA* mRNA expression level was increased in WFS1-deficient β-cells. VEGF-A produced in pancreatic islets plays a critical role in islet angiogenesis (26), and we have previously shown that VEGF-A transcription is regulated by the UPR pathway (35). Islet-resident macrophages and blood vessels are in close contact, and chronic islet hypervascularization leads to progressive macrophage infiltration (25, 36). WFS1-deficient cells are vulnerable to ER stress, which promotes *VegfA* expression, which may lead to hypervascularization and macrophage infiltration in the WS model mice islets. Further studies are needed to clarify whether intra-islet macrophages are proliferating or whether monocytes are recruited into the islets.

A high-glucose environment is known to induce pro-inflammatory cytokine and chemokine genes in macrophages (37). However, in this study, we show that the expression levels of pro-inflammatory cytokine genes in high-glucose treated bone marrow-derived macrophages were not altered by WFS1 deficiency. These results indicate that the functional contribution of WFS1 in the macrophages is not as significant as in the pancreatic β-cells, possibly because the expression level of WFS1 is lower in macrophages than in the other organs (https://www.proteinatlas.org/ENSG00000109501-WFS1). On the other hand, *Wfs1*-deficient bone marrow-derived macrophages stimulated with AGE showed an increased expression level of the *Ccl2* gene, a marker of M1 macrophages. Since AGE serum levels in Wolfram syndrome model mice increased with age, it is possible that stimulating factors other than blood glucose, including AGE, alter the transcript signature of macrophages in a model of Wolfram syndrome.

In type 2 diabetes, elevated levels of serum pro-inflammatory cytokines such as IL-1β and IL-6 are predictive disease markers (14, 38, 39). In this study, we showed that inflammatory cytokine levels in the serum of a mouse model of Wolfram syndrome are higher than that of wild type mice. This result is consistent with the recent studies that showed elevated cytokine levels in WS patient serum and elevated cytokine production in peripheral blood mononuclear cells (PBMC) isolated from Wolfram syndrome patients (40, 41). The source of the elevated serum cytokines in Wolfram syndrome is still unknown; however, since the pancreatic islets are highly vascularized, systemic inflammation may affect the islet inflammation. It is essential to investigate the contribution to the systematic inflammation in Wolfram syndrome by other tissues and organs such as adipose tissue, liver, and blood cells other than macrophages, which are involved in the inflammation associated with type 2 diabetes. Cells or tissues differentiated from patient-derived iPSCs would provide a better model to study the role of inflammation in Wolfram syndrome (42).

Besides the Wolfram syndrome β-cells, sterile inflammation occurs in many other cell types such as oligodendrocytes, hepatocytes, and adipocytes, where the ER is heavily loaded with proteins (9). Symptoms seen in Wolfram syndrome are accompanied not only by diabetes but also by optic nerve atrophy and neurodegeneration, whose pathogenic mechanisms are not fully understood. Therefore, this study suggests that the metabolic factors, including hyperglycemia, inflammatory cytokines, and AGE, may contribute to the progression of Wolfram syndrome symptoms in a cell-nonautonomous manner.

In summary, our study identifies sterile inflammation as a new pathological mechanism in the Wolfram syndrome. In Wolfram syndrome model mice, islets are highly infiltrated with macrophages, suggesting that the vicious cycle of ER stress and inflammation accelerates the progression of diabetes in a cell-autonomous and cell-nonautonomous manner. A deeper understanding of the pathophysiology of Wolfram syndrome is important for developing novel therapeutic targets for the disease. As the inflammation-targeted therapies have been tried in type 2 diabetes (43), the findings from this study may also be applicable not only to diabetes in Wolfram syndrome but also to other manifestations such as optic atrophy and neurodegeneration, which severely impair the quality of life of patients with Wolfram syndrome.

## Materials and Methods

### Cell culture

*Wfs1*-KO INS-1 832/13 cells were generated in collaboration with Genome Engineering and Induced Pluripotent Stem Cell Center (GEiC) at Washington University in St. Louis as described before (17). INS-1 832/13 cells were cultured in 11 mM glucose RPMI 1640 (Thermo Fisher Scientific, Cat# 11875) supplemented with 10% fetal bovine serum (FBS) (Thermo Fisher Scientific, Cat# 10099141), 1 mM sodium pyruvate (Corning, Cat# 25-000-CI), 90 µM β-mercaptoethanol (MilliporeSigma, Cat# M3148) and 100 U/mL penicillin-streptomycin (Thermo Fisher Scientific, Cat# 15140122). INS-1E cells were cultured in 11 mM RPMI 1640 supplemented with 10% FBS, 1 mM sodium pyruvate, 50 µM β-mercaptoethanol, 2 mM GlutaMAX (Thermo Fisher Scientific, Cat# 35050061), NEAA (Thermo Fisher Scientific, Cat# 11140050), 10 mM HEPES (Corning, Cat# 25-060-CI), and 100 U/mL penicillin-streptomycin. Rat IFN-γ, rat IL-1β (R&D Systems, Cat# 585-IF-100, 501-RL-010), and thapsigargin (MilliporeSigma, Cat# T9033) were used for the treatments.

### Wolfram syndrome mouse model and pancreatic sample preparation

129S6 whole body *Wfs1*-knockout mice were a kind gift from Dr. Sulev Kõks (University of Tartu) (22). Amino acids 360-890 of WFS1 protein were replaced with an in□frame NLSLacZNeo cassette. Mouse genotypes were determined using multiplex PCR performed by Transnetyx (Cordova, TN). All animal experiments were performed according to procedures approved by the Institutional Animal Care and Use Committee at the Washington University School of Medicine (Protocol # 20-0334). All mice were housed in a pathogen-free animal facility, and food and water were provided *ad libitum* throughout the study. For the pancreatic section slides, 129S6 whole body *Wfs1*-knockout mice were euthanized in a carbon dioxide chamber followed by cervical dislocation. The mice were perfused with PBS and 4% paraformaldehyde (PFA) via the left ventricle. After perfusion, the pancreas was removed and fixed again with 4% PFA for 48 h at 4°C. The fixed pancreas was gradually replaced by ethanol and finally kept in 70% ethanol. Paraffin embedding and sectioning were performed by Histology and Morphology Core at Musculoskeletal Research Center at Washington University in St. Louis. Slides were sectioned at 5 µm thickness. Microscopy imaging was performed using a Leica DM6B (Leica Microsystems, Germany). All quantitative analyses were conducted using Fiji ImageJ in a double-blinded fashion.

### Immunostaining

Paraffin-embedded pancreatic section slides were deparaffinized, and antigen retrieval was performed by submerging the slides in 10 mM sodium citrate solution (pH 6.0) for 30 min at 95 °C. The tissues were permeabilized with phosphate buffer saline (PBS) containing 0.1% Triton X-100 for 30 min at room temperature. For the DAB stained samples, endogenous peroxidase was quenched using BLOXALL Blocking solution (VECTOR, Cat# SP-6000). Blocking was performed in 2% bovine serum albumin (BSA) for 1 h. The following primary antibodies were diluted in 0.2% BSA and incubated overnight at 4 °C: anti-WFS1 (Proteintech, Cat# 1158-1-AP, 1:100), anti-Iba1 (Novus, Cat# NB100-1028, 1:50), anti-CD31 (abcam, Cat# ab124432, 1:100), and Alexa-Fluor 488 conjugated anti-insulin (Invitrogen, Cat# 53-9769-82, 1:100). After washing the primary antibodies 3 times with PBS, the tissues were incubated for 30 min at room temperature with ImmPRESS polymer reagent (VECTOR, Cat# MP-7401-50) for DAB staining, or AlexaFluor 594 donkey anti-rabbit IgG (Invitrogen, Cat# A21207) for immunofluorescent staining, and washed for 2 times with PBS. The color development for DAB staining was performed using peroxidase substrate solution (VECTOR, Cat# SK-4100) following the manufacturer’s protocol, and then the nucleus was stained by Vector Hematoxylin QS (VECTOR, Cat# H-3404). Tissues were mounted in Vectamount permanent mounting medium (VECTOR, Cat# H5000-60).

### Masson’s trichrome staining

Deparaffinized and rehydrated slides were immersed in Bouin’s solution at 56°C for 1 h. Subsequently, the slides were washed with tap water for 5 min. The washed sections were stained in Weigert’s Hematoxylin solution for 10 min and then washed again with tap water for 5 min. Next, the slides were stained in Trichrome solution for 15 min and rinsed in 1% acetic acid for 1 min. Slides were dehydrated in alcohol twice for 1 min each, and sections were cleared in xylene twice for 1 min each. All the Masson’s trichrome staining procedures were performed by the Musculoskeletal Research Center at Washington University in St. Louis.

### Imaging mass cytometry

The antibody staining on formalin-fixed paraffin-embedded (FFPE) mouse pancreatic tissue was performed based on Fluidigm immunohistochemistry protocol. Briefly, the FFPE slides were deparaffinized followed by antigen retrieval using 10 mM sodium citrate (pH 6.0) for 30 min at 96°C. Blocking was performed using metal-free 3% BSA for 45 min prior to the addition of the primary antibody solution. The primary antibodies were anti-F4/80(BM8) (Fluidigm, Cat# 3146008B), anti-CD68(FA-11) (Biolegend, Cat# 137001), anti-CD163(EPR19518) (Abcam, Cat# ab182422), and anti-IL-1β/IL-1F2(Novus Biologicals, Cat# NBP1-42767). Metal conjugation was performed using MAXPAR® X8 Multimetal Labeling Kit (Fluidigm, Cat# 201300), and all the antibodies were used in a 1:25 dilution. After overnight incubation of the antibody mixture at 4°C in a humidity chamber, the slide was washed with 0.2% Triton X-100 in PBS and then stained with 0.0025% Ruthenium Red for contrast staining. After the contrast staining, DNA intercalator for nuclear identification was added. The slides were washed in distilled water and then air-dried before imaging. Antibody conjugation and CyTOF2/Helios imaging were performed by the Immunomonitoring Laboratory (IML) in Bursky Center for Human Immunology & Immunotherapy Programs (CHiiPs) at Washington University in St. Louis. Optimization of the multiplex panel involves assessing dual signal spillover into +1, +2, and +16 channels and lack of signal (false negative). Assessment of signal in other channels was performed using MCD Viewer software and generating a thumbnail image file for each metal channel. The imaging data were converted to TIFF images using HistoCAT++ (ver 3.0.0), and CellProfiler (ver 4.1.3) was used for the cell masking. Image visualization and data analysis were performed using HistoCAT (ver 1.761).

### Cell death assay

INS-1 832/13 cells were plated on 96-well flat clear bottom white polystyrene plates (Corning, Cat# 3610) and treated with or without the indicated reagents. After the treatment, CellTiter-Fluor Cell Viability Assay (Promega, Cat# G6080) reagent was added directly to cells and incubated for 30 min in the dark. After the fluorescence measurement, Caspase-Glo 3/7 Assay (Promega, Cat# G8090) reagent was added to the cells and incubated for 30 min. Fluorescence for cell viability and luminescence for caspase-3/7 activity was measured using Infinite M1000 plate reader (Tecan). Caspase-3/7 activity was normalized to cell viability according to the manufacturer’s protocol.

### Immunoblot

Cells were washed in cold PBS and immediately lysed on ice in Mammalian Protein Extraction Reagent (Thermo Fisher Scientific, Cat# 78501) supplemented with 1X cOmplete^**™**^ protease inhibitor cocktail (MilliporeSigma, Cat# 11873580001) and 1X PhosSTOP^**™**^ phosphatase inhibitor (MilliporeSigma, Cat# 4906845001) before centrifugation at 15,000 rpm for 10 min at 4°C. Nuclear and cytoplasmic extraction was performed using NE-PER™ (Thermo Fisher Scientific, Cat# 78833) according to the manufacturer’s protocol. Protein lysates were prepared using 4x Laemmli sample buffer (Bio-Rad Laboratories, Cat# 1610747) heated at 60°C for 15 min. The protein was resolved by SDS-PAGE and transferred to Immobilon-P PVDF membrane (MilliporeSigma, 0.2 µm pore size (Cat# ISEQ20200) for proinsulin and 0.45 µm pore size for others (Cat# IPVH00010)). For detecting proinsulin, additional fixation and antigen retrieval steps were performed as reported previously (44). The antibodies used for immunoblotting were anti-WFS1 (Proteintech, Cat# 1158-1-AP), anti-LaminB1 (Proteintech, Cat# 12987-1-AP), and those purchased from Cell Signaling Technology: anti-cleaved caspase-3 (Cat# 9664), anti-insulin (Cat# 8138), anti-pPERK (Cat# 3179), anti-PERK (Cat# 3192), anti-IRE1α (Cat# 32945), anti-NF-κB (Cat# 8242), anti-GAPDH (Cat# 2118), anti-α-tubulin (Cat# 2125), and the secondary antibodies conjugated to horseradish peroxidase. Bands were detected by ECL Select (MilliporeSigma, Cat# RPN2235) and Bio-Rad ChemiDoc MP. Quantitative analyses were conducted using Fiji ImageJ.

### Quantitative real-time PCR

Total RNA was extracted from INS-1E, INS-1 832/13, or isolated primary mouse macrophages using RNeasy Mini Kit (Qiagen, Cat# 74106) and reverse-transcribed using High-Capacity cDNA Reverse Transcription Kits (Thermo Fisher Scientific, Cat# 4368814). The qPCR was performed in 3-8 replicates for each sample. All the qPCR primer sequences used in this study are listed in Table S1.

### siRNA

INS-1E or INS-1 832/13 cells were seeded in a 12-well plate and transfected with siRNA using RNAiMAX (Thermo Fisher Scientific, Cat# 13778150). ON-TARGET plus SMARTpool siRNA was used for knocking down rat *Wfs1* (Dharmacon, Cat# L-087932-02). The siRNA against rat *Perk, Ire1*α, and non-targeting siRNA were purchased from MilliporeSigma (rat *Perk*, SASI_Rn01_00064452; rat *Ire1*α, SASI_Rn02_00372535; SIC001 for non-targeting). Each siRNA was used in 40 nM concentration, and RNAiMAX was used 1.0 µL against 1 pmol of siRNA. The cells were collected for the analysis 48 h after the siRNA transfection.

### Statistical analysis

All statistics were performed using GraphPad Prism 9. Statistical analysis was performed by unpaired two-tailed Student’s t-test or one-way ANOVA followed by Tukey’s multiple comparisons test. P <0.05 was considered statistically significant.

## Acknowledgment

F. Urano thanks philanthropic supports from the Silberman Fund, the Ellie White Foundation for the Rare Genetic Disorders, the Snow Foundation, the Unravel Wolfram Syndrome Fund, the Stowe Fund, the Eye Hope Foundation, the Feiock Fund, Ontario Wolfram League, Associazione Gentian - Sindrome di Wolfram Italia, Alianza de Familias Afectadas por el Sindrome Wolfram Spain, Wolfram syndrome UK, and Association Syndrome de Wolfram France. F. Urano also thanks all the members of the Washington University Wolfram Syndrome Study, Research Clinic, and WFS1 Clinic at the Washington University Medical Center for their support (https://wolframsyndrome.wustl.edu) and all the participants in the Wolfram syndrome International Registry and Clinical Study, Research Clinic, and Clinical Trials for their time and efforts We acknowledge Diane Bender and Kohei Omachi (both in Washington University in St. Louis) for their skilled technical supports and Kohsuke Kanekura (Tokyo Medical University) for his critical review of the manuscript. The authors are grateful to Cris Brown (Washington University in St. Louis) for her general support. This work was supported, in part, by the Bursky Center for Human Immunology and Immunotherapy Programs at Washington University in St. Louis, Immunomonitoring Laboratory. We are grateful for the critical review and editing assistance provided by InPrint: A Scientific Communication Network at Washington University in St. Louis.

## Author contribution

S.M. and F.U. conceived the project. S.M. designed the experiments. S.M. and L.B. performed the experiments, and S.M., L.B., and C.O. analyzed the data. F.U. supervised the work. All authors participated in writing, editing, and reviewing the manuscript.

