## Supplemental Data for "Anti-inflammatory effects of WFS1 in pancreatic β-cells"

**Supplemental Information**

### Materials and Methods

#### Primary mouse macrophage isolation

Eight-week-old mice were euthanized, and their hind legs were cleaned with 70 % ethanol. Femur and tibia bones were harvested by cutting off the hind legs close to the hip joint. The femur and tibia bones were wiped with 70% ethanol, and both ends were cut. The bones were placed into PCR tubes with a needle hole on the bottom. The PCR tubes were placed in a 1.5 mL tubes and filled with phosphate buffer saline (PBS). After centrifuging at 13,000 rpm for 10 min at 4°C, the pellets were suspended in 10 mL PBS and filtered with 70 µm cell strainers. The flowthrough was centrifuged at 1,000 rpm for 10 min at 4°C, and the pellets were resuspended with 4 mL warm macrophage culture medium (MEM-alpha (5.5 mM glucose, Thermo Fisher Scientific, Cat# 12561049) containing 10% fetal bovine serum (FBS), 100 U/mL penicillin-streptomycin, and 50 ng/mL M-CSF (STEMCELL, Cat# 78059)). The suspended cells were seeded on 6-cm petri-dishes and incubated at 37°C in 5% $CO_2$. After overnight incubation, the cells were plated on 12-well plates. The cells were cultured in macrophage culture media for 7 days. Half of the media was replaced every other day. On day 7, the cells were stimulated with 5.5 mM or 30 mM glucose for 24 h, or 200 µg/mL BSA or Advanced Glycation Endproduct-BSA (MilliporeSigma, Cat# 121800-M) for 6 h.

#### Mitochondrial function assay

Mitochondrial function was assessed by measuring the oxygen consumption rate (OCR) using a Seahorse XFe96 analyzer (Agilent Technologies). *Wfs1* wild type INS-1 832/13 (*Wfs1*-WT INS-1) and knock-out INS-1 832/13 (*Wfs1*-KO INS-1) cells were seeded on Seahorse 96-well cell culture plate (Agilent Technologies, Cat# 101085-004) before the day of the assay. On the day of the assay, the media was replaced with Agilent Seahorse XF RPMI with 1 mM HEPES (Agilent Technologies, Cat# 103576-100) supplemented with 1 mM Seahorse XF Pyruvate (Agilent Technologies, Cat# 103578-100), 2 mM Seahorse XF Glutamine (Agilent Technologies, Cat# 103579-100), and 11 mM Seahorse XF Glucose (Agilent Technologies, Cat# 103577-100). The plate was placed in a non-$CO_2$ incubator at 37°C for 1 h, and then placed in a Seahorse XFe96 Analyzer. The concentration of the electron transport chain inhibitors and uncouplers were oligomycin (1.5 µM), carbonyl cyande-4-(trifluoromethoxy) phenylhydrazone (FCCP, 1.0 µM), rotenone (1.0 µM), and antimycin A (1.0 µM). All compounds were purchased from MilliporeSigma. Four OCR measurements were recorded for baseline and following each compound injection. The OCR parameters were calculated by subtracting the average respiration rates before and after the addition of the electron transport inhibitors.

#### Intraperitoneal glucose tolerance test

Ten-month-old male 129S6 wild type or whole body *Wfs1*-knockout mice were used for intraperitoneal glucose tolerance test (IPGTT). 50% dextrose (Hospira, NDC 0409-6648) was used for the intraperitoneal injection. The animal number for each genotype group is indicated in the figure legend. The test was performed according to standard procedures of the NIH-sponsored National Mouse Metabolic Phenotyping Centers (http://www.mmpc.org).

**Mouse serum samples**

Mouse blood samples were collected from the tail vein. After the whole blood collection and coagulation, the clot was removed by centrifuging at 2,000 g for 10 min at 4°C. Mouse serum advanced glycation end-product (AGE) was measured by OxiSelect Advanced Glycation End Product (AGE) Competitive ELISA kit (Cell Biolabs, Cat# STA-817) using a Bio-Tek ELx800 Plate reader. The assay was performed by Core Laboratory for Clinical Studies (CLCS) at Washington University in St. Louis.

**Fig. S1**

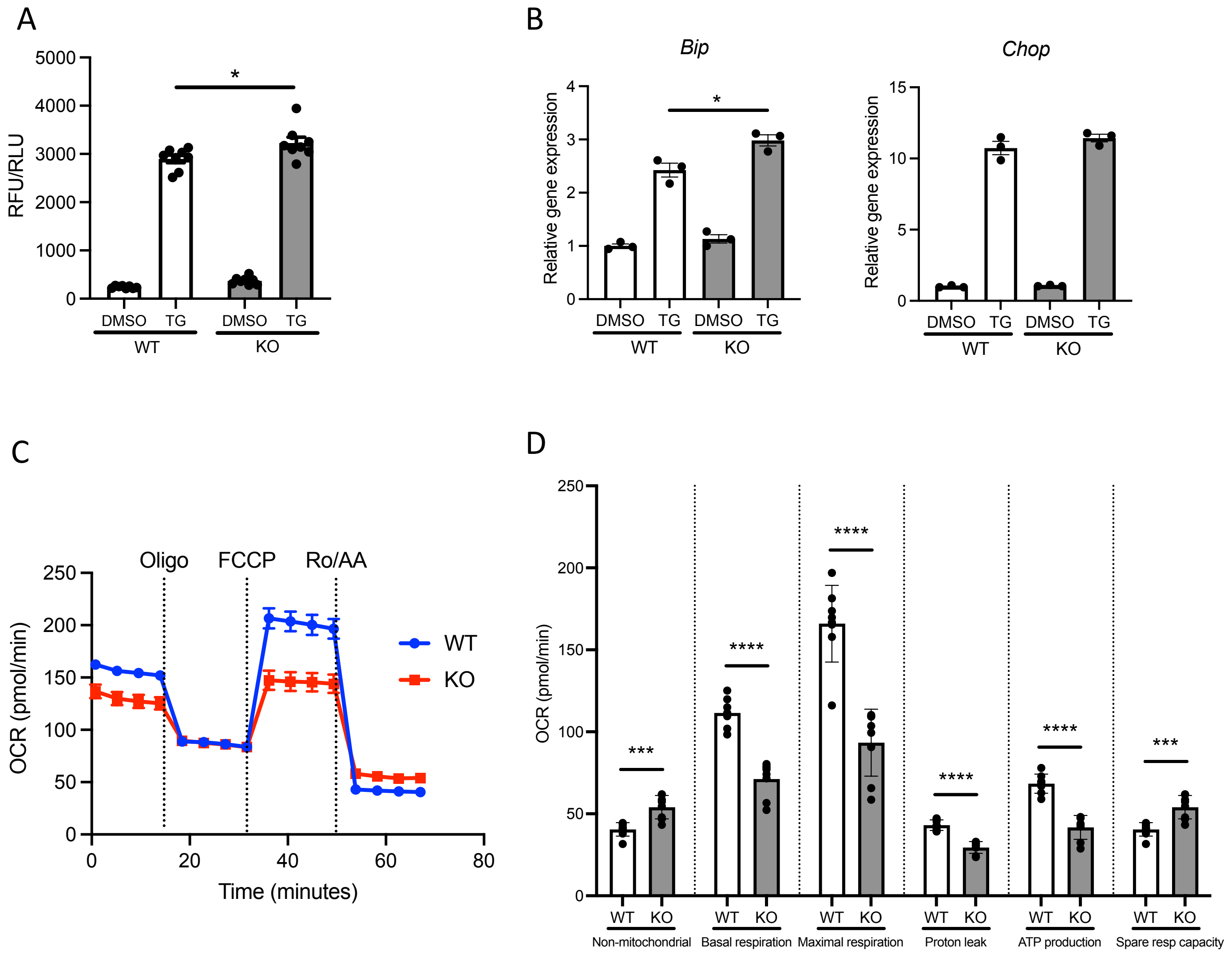


**Fig.S1 ER stress intolerance and mitochondrial dysfunction in *Wfs1*-KO INS-1 832/13 cells**

(A) Caspase-3/7 activity normalized to cell viability in *Wfs1* wild type (WT) and knock-out (KO) INS-1 832/13 cells treated with DMSO or thapsigargin (TG, 100 nM) for 4 h (n=8). (B) mRNA expression levels of *Bip* and *Chop* in WT or KO INS-1 832/13 cells treated with DMSO or thapsigargin (TG, 100 nM) for 4 h (normalized to *18srRNA*, n=3). (C) Oxygen consumption rate (OCR) in WT or KO INS-1 832/13 cells. Oligo, oligomycin; FCCP, carbonyl cyande-4-(trifluoromethoxy) phenylhydrazone; Ro, rotenone; AA, antimycin A. (D) Calculated OCR parameters in WT or KO INS-1 832/13 cells (n=8). Data are shown as mean ± SEM, * P<0.05, *** P<0.001, **** P<0.0001.

**Fig. S2**

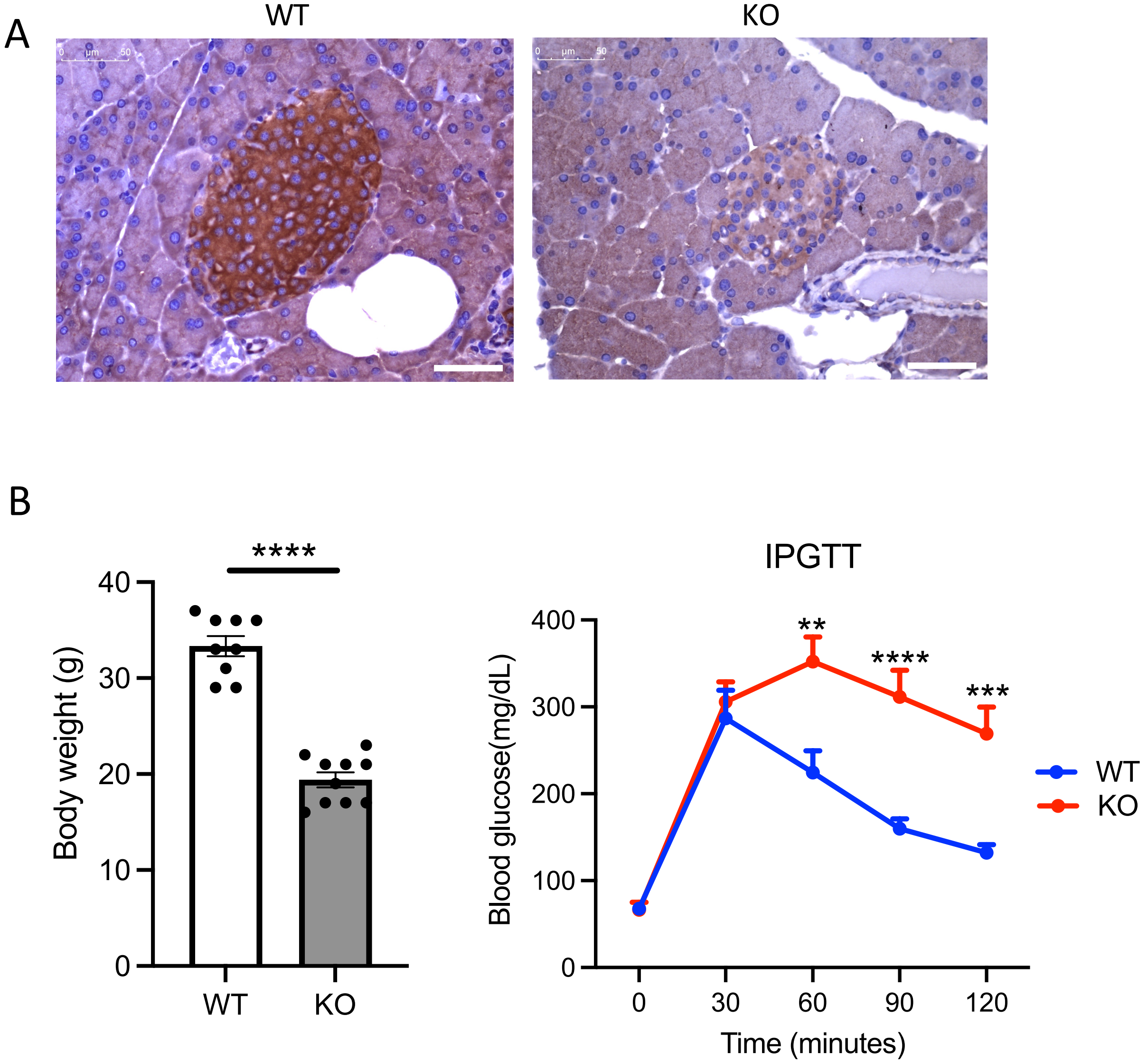


**Fig.S2 Whole body Wfs1$^{-/-}$ mice show the diabetic symptoms**

(A) WFS1 localization in *Wfs1* wild type (WT) and whole-body knockout 129S6 (KO) mouse islet. WFS1 staining of the islets from WT and KO male mice at 17 weeks of age. High expression of WFS1 was observed in WT mice islets. In contrast, no WFS1 protein expression and disrupted or small islet architecture were observed in KO mice. Scale bars 50 µm. (B) Left panel: Body weight in 10-month-old WT and KO mice (unpaired two-tailed Student's t-test, **** P<0.0001). Right panel: Intraperitoneal glucose tolerance test (IPGTT) results in 10-month-old WT and KO mice (WT, n=9; KO, n=10, two-way ANOVA, ** P<0.005, *** P<0.001, **** P<0.0001). Data are shown as mean ± SEM.

**Fig. S3**

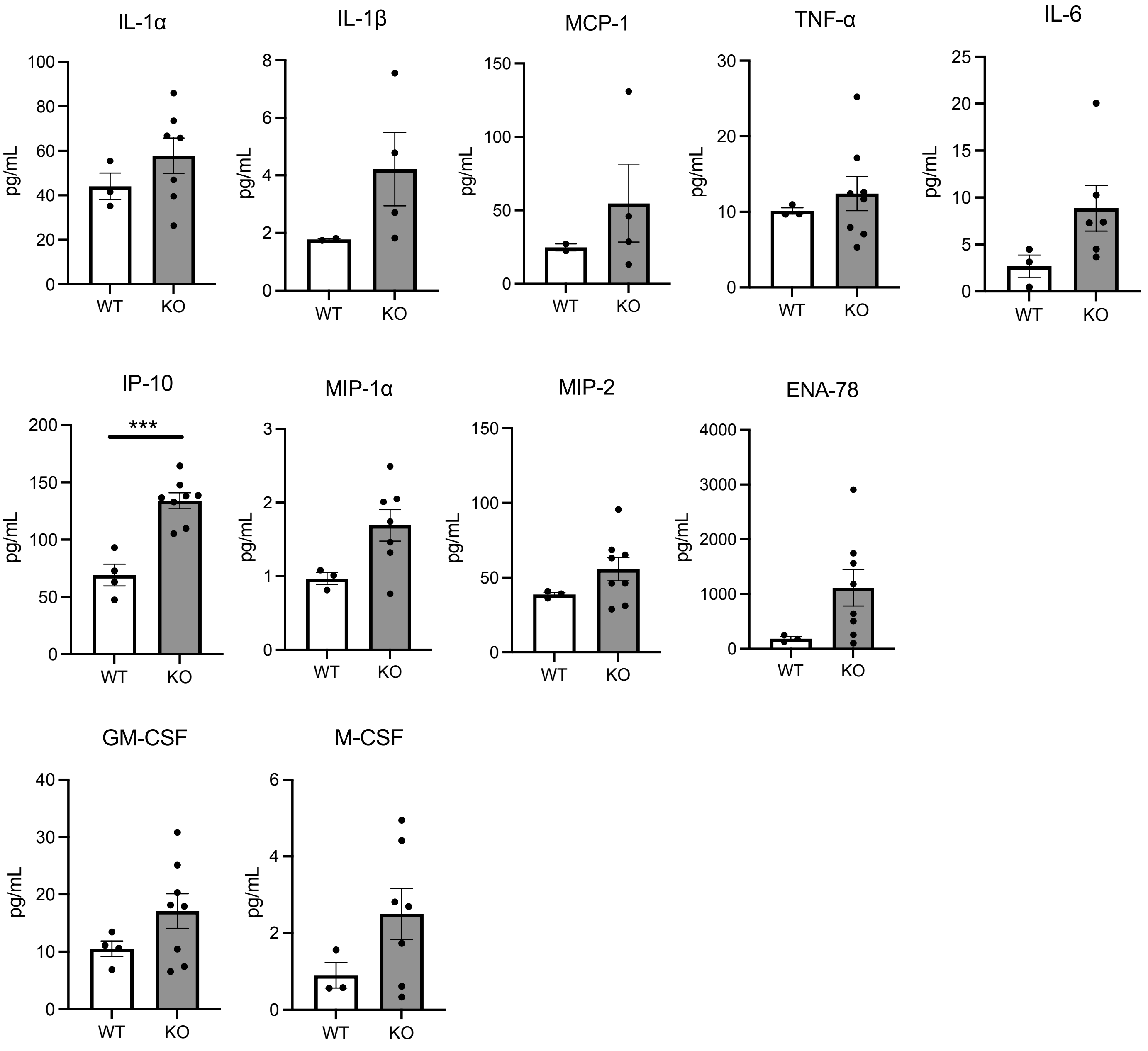


**Fig.S3 Systematic inflammation in Wolfram syndrome mouse model**

Serum cytokine levels in 5-month-old *Wfs1* wild type (WT) and whole-body knockout 129S6 (KO) female mice (n=3-8). Data are shown as mean ± SEM, *** P<0.001.

**Fig. S4**

**Mouse bone marrow derived macrophages**

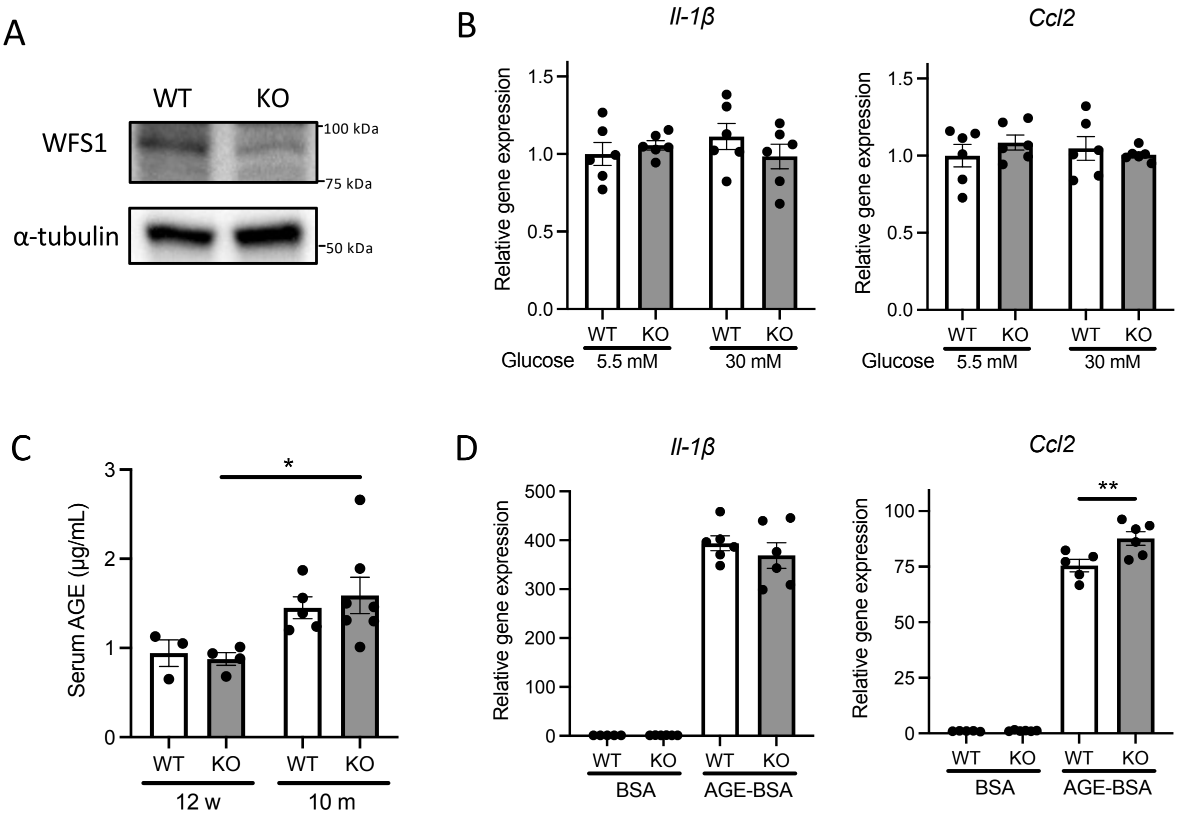


**Fig.S4 Advanced glycation end-products (AGE) upregulates the gene expression level of Ccl2 in Wfs1-deficient bone marrow delivered macrophages.**

(A) Immunoblot images of WFS1 in primary bone marrow delivered macrophages (BMDM) isolated from 7-week-old *Wfs1* wild type (WT) and whole-body knockout 129S6 (KO) male mice. (B) mRNA expression levels of *Il-1β* and *Ccl2* in BMDM treated with 5.5 mM or 30 mM glucose for 24 h. mRNA levels are normalized to *Gapdh*, n=6. (C) Serum level of advanced glycation end-product (AGE) in WT and KO male mice at 12-week-old and 10-month-old (n=3-7). (E) mRNA expression levels of *Il-1β* and *Ccl2* in BMDM treated with bovine serum albumin (BSA) or AGE-BSA (200 µg/mL) for 6 h (n=5-6). mRNA levels are normalized to *Gapdh*. Data are shown as mean ± SEM, * $P<0.05$, ** $P<0.005$

**Table S1 The qPCR primer sequences used in this study**

| Species | Gene name | Forward Primer Sequence | Reverse Primer Sequence |
|---|---|---|---|
| Rat | *sXbp1* | CTGAGTCCGAATCAGGTGCAG | ATCCATGGGAAGATGTTCTGG |
| | *Bip* | TGGGTACATTTGATCTGACTGGA | CTCAAAGGTGACTTCAATCTGGG |
| | *Chop* | AGAGTGGTCAGTGCGCAGC | CTCATTCTCCTGCTCCTTCTCC |
| | *Txnip* | CAAGTTCGGCTTTGAGCTTC | ACGATCGAGAAAAGCCTTCA |
| | *Il-1β* | CTCTGTGACTCGTGGGATGA | CGAGGCATTTTTGTTGTTCA |
| | *Il-6* | TAGTCCTTCCTACCCCAACTTCC | TTGGTCCTTAGCCACTCCTTC |
| | *Cxcl1* | CATTAATATTTAACGATGTGGATGCGTTTCA | GCCTACCATCTTTAAACTGCACAAT |
| | *Ccl2* | CTTCTGGGCCTGTTGTTCAC | GCCAGTGAATGAGTAGCAGC |
| | *VegfA* | CAAGCCGTCCTGTGTGCC | TCCAGGGCTTCATCATTGC |
| | *18srRNA* | GCAATTATTCCCCATGAACG | GGCCTCACTAAACCATCCAA |
| Mouse | *Il-1β* | TGGACCTTCCAGGATGAGGACA | GTTCATCTCGGAGCCTGTAGTG |
| | *Ccl2* | GCTACAAGAGGATCACCAGCAG | GTCTGGACCCATTCCTTCTTGG |
| | *Gapdh* | TGTAGACCATGTAGTTGAGGTCA | AGGTCGGTGTGAACGGATTTG |
